# Echolocating bats prefer a high risk-high gain foraging strategy to increase prey profitability

**DOI:** 10.1101/2022.10.25.513681

**Authors:** Laura Stidsholt, Antoniya Hubancheva, Stefan Greif, Holger R. Goerlitz, Mark Johnson, Yossi Yovel, Peter T. Madsen

## Abstract

Most bats catch nocturnal prey during active flight guided by echolocation but some species depart from this ancestral behaviour to capture ground prey using passive listening. Here, we explore the costs and benefits of these hunting transitions by combining high-resolution biologging data and DNA metabarcoding to quantify the relative contributions of aerial and ground prey to the total food intake of wild greater mouse-eared bats. We show that these bats use both foraging strategies with similar average nightly captures of 25 small, aerial insects and 30 large, ground-dwelling insects per bat, but with higher capture success in air (78 % in air vs 30 % on ground). However, owing to the 3 to 20 times heavier ground prey, 85 % of the estimated nightly food acquisition comes from ground prey despite the 2.5 times higher failure rates. Further, we find that most bats use the same foraging strategy on a given night suggesting that bats adapt their hunting behaviour to weather and ground conditions. We conclude that prey switching matched to environmental dynamics plays a key role in covering the energy intake even in specialised predators.

## Introduction

Bats are widespread and abundant predators that serve important roles in ecosystems across the globe (Kunz *et al*., 2011).Their evolutionary success is due to the unique combination of echolocation and powered flight (Teeling *et al*., 2005) allowing them to avoid visual predators by feeding at night, thereby gaining unfettered access to food sources that include insects, small vertebrates, fruit, nectar, pollen and blood(Simmons, 2005). Within the mosaic of foraging niches they exploit, echolocating bats are categorised into three main foraging strategies: the ancestral mode of capturing prey on the wing (hawking), and the derived modes of trawling prey from water surfaces or gleaning prey, nectar or fruit from ground and trees(Schnitzler and Kalko, 2001). To aid these specialised hunting strategies, each guild of bats has evolved specific adaptations in echolocation signals, auditory systems, morphology, and flight mechanics (Norberg and Rayner, 1987; Fenton, 1990; Schnitzler and Kalko, 2001). Despite such specialism, recent research has shown that foraging style is not monotypic within species: gleaning bats occasionally capture aerial prey (Bell, 1982; Fenton, 1990; Ratcliffe and Dawson, 2003; Ratcliffe, Fenton and Shettleworth, 2006; Hackett, Korine and Holderied, 2014), while insect-gleaning bats may seasonally target nectar or fruit(Aliperti *et al*., 2017) or vice versa (Herrera *et al*., 2001). These changes in foraging style presumably track the relative abundance of preferred versus alternative food sources, broadening the ecological roles of bats and providing a degree of resilience in the face of changing resources. However, owing to the complexity of studying detailed hunting behaviours in the wild, it is not clear why or when specialised bats switch foraging strategies. This led us to ask whether bats adapt their hunting strategies continuously to maintain net intake or if switching is the last resort when preferred prey are unavailable. To address that, we used miniaturized biologging devices to track the hunting behaviour of greater mouse-eared bats. This species primarily captures ground-dwelling arthropods by passively listening for their movements(Arlettaz, 1996), and is therefore specialised for gleaning: broad, short wings enable them take-off from the ground, and weak echolocation calls avoid alerting prey while also allowing the bat to hear rustling sounds of surface-dwelling prey. Nonetheless, like many other gleaning bats, greater mouse-eared bats have maintained the ancestral ability to use echolocation to capture aerial prey on the wing, which requires that they switch to intense calls to detect small prey, and maintain the capability to manoeuvre fast in 3D space to track evasive prey. While call intensity can be adjusted to fit different strategies, the morphological and anatomical specialisations for ground gleaning must affect the efficiency of these bats as aerial hawkers. We therefore predicted that bats would prefer gleaning whenever it was profitable, and would only switch to aerial hunting when environmental conditions led to poor prey biomass per foraging time when gleaning. To test this prediction, we used sound and movement tags (N = 34 bats) (Stidsholt *et al*., 2018) to record the bats’ echolocation behaviour, 3D movement patterns, GPS locations (N = 7 of 34 bats) and mastication sounds after prey captures to quantify foraging success. We augmented these data with DNA metabarcoding of faeces from co-dwelling con-specifics (N = 54 bats) to identify prey species and sizes. This combined biologging and DNA metabarcoding approach allows us to estimate prey profitability across foraging strategies and habitats to evaluate the drivers of prey switching in wild bats.

## Results

To categorise the foraging strategies used by wild bats, we analysed one night of sound and movement data from each of 34 female, greater mouse-eared bats (*Myotis myotis*). All bats commuted from the colony or release site to one or several foraging grounds before returning back to the roost before dawn (Fig. 1ABC). Since we used two different types of biologging devices with different sensors, foraging bouts were defined either as intervals of high variation in heading, or as intervals of area-restricted search for tags including GPS (N = 7 bats). A total of 3917 attacks on prey were recorded with most bats capturing prey both on the ground by passively gleaning prey, and by pursuing prey mid-air by aerial hawking (Fig. 1DE). However, four bats exclusively gleaned, while two bats only hawked (Fig. 1DE). The dominant foraging strategy used per bat per night seemed to be affected by the night of tagging indicating that bats tagged on the same nights choose the same foraging strategy (Fig. 1D & Fig. S1, N = 10 nights, 1 to 9 bats tagged per night; LMM; testing if the ratio between ground:aerial captures was explained by night of tagging, p=0.001).

**Fig. 1:**
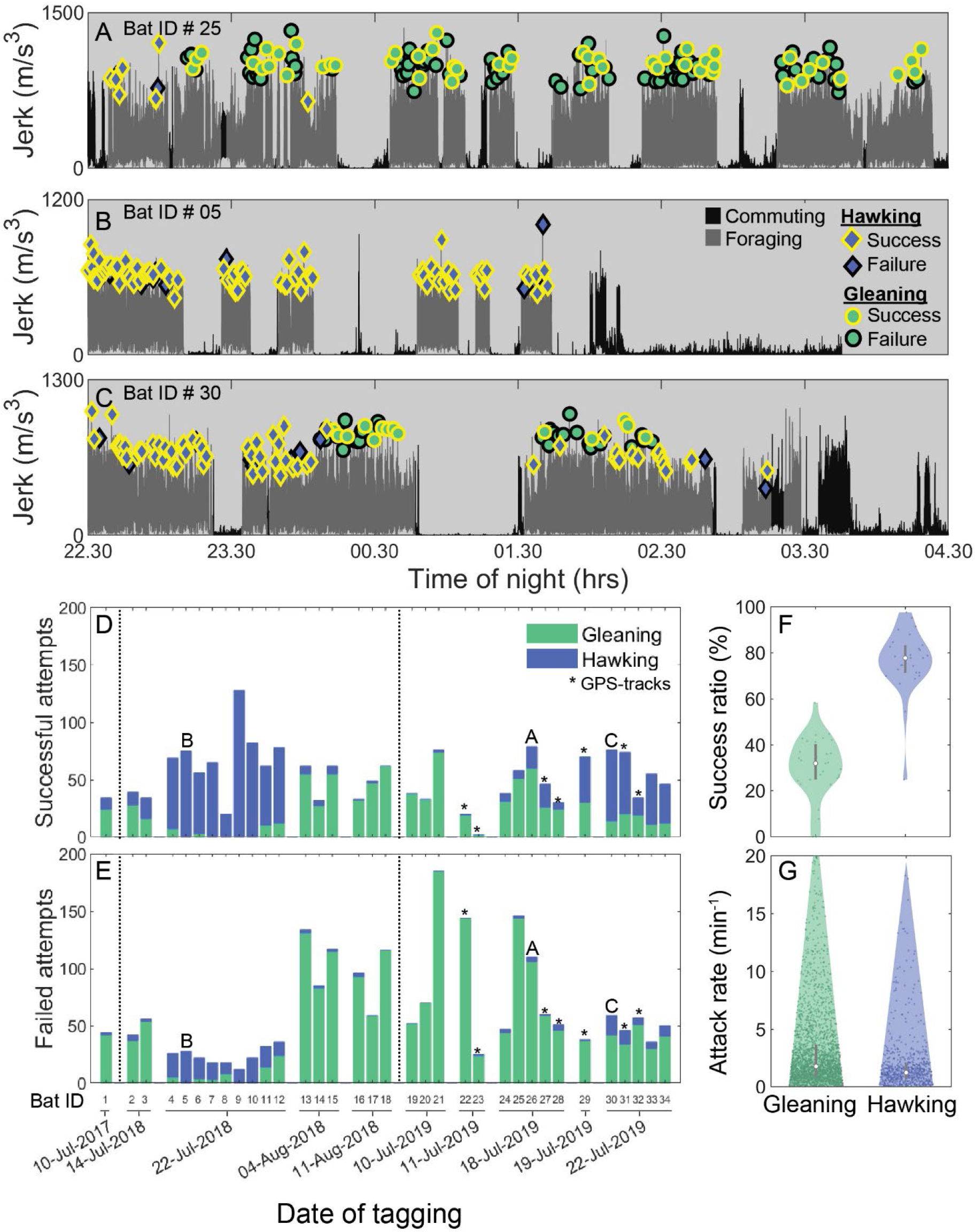
Greater mouse-eared bats tagged on different nights show wide variation in foraging strategy and success. A-C: The jerk (differential of on-animal recorded acceleration) reveals the overall movement of the bat by showing periods of no movement (rest) and strong movement (flight) for three different bats (summed values for Bat ID 25,5 and 30 as depicted in panel D) and two different travel modes (commuting (dark grey) vs foraging (light grey)). We marked all prey attacks as either hawking (blue diamonds) or gleaning (green circles) by visual and auditory inspection of the sound and movement data. Prey captures were classified by audible mastication sounds as successful (black edge) or failures (pink edge). The bats exemplified here either primarily gleaned (A), primarily hawked (B) or used both strategies in alternating bouts (C). D-E) Successful (D) and unsuccessful (E) prey attacks of all bats (N = 34) grouped according to night of tagging for aerial hawking (blue) and gleaning (green). Stars mark the bats equipped with GPS tags; A, B, C mark the bats depicted in panels A-C. F-G) The success ratio (F) reveals the percentage of all attacks that were successful per bat per night (dots), while attack rates (G) reveal the number of foraging attacks per minute for each bat per night (dots) with more than one prey attack per foraging strategy for aerial hawking (blue) and gleaning (green) along with kernel densities and boxplots.

The bats attacked food more often on the ground (mean: 80 attacks per individual per night, quartiles: 26–110) than in the air (mean: 35 attacks, quartiles: 6–70; LMM, p=0, Table S3, Fig. 1D), but the proportion of attacks that were successful (*i.e*. success ratio) based on audible mastication sounds following prey captures were more than double in air (79 %, quartiles: 71– 88) than on ground (31 %, quartiles: 25–40; Table S4, Fig. 1F). This led to on average 25 (quartiles: 10 to 33) aerial and 30 (quartiles: 5 to 53) ground insects caught per bat per night (Fig. 1D). Prey attack rates were substantially higher for ground foraging versus aerial foraging (ground: 1.73 (quartiles: 1.7-1.75) prey attacks/minute vs aerial: 1.17 (quartiles: 1.14-1.21)) (Fig. 1G). Thus, bats caught prey much more reliably in the air, but attacked ground prey more often and devoted more foraging time to ground gleaning.

We next used GPS tracks from seven bats to investigate the behavioural and ecological factors that influence foraging success. Specifically, we tested if movement style (i.e., commuting vs actively searching for prey defined via the Lavielle method(Hurme *et al*., 2019)) and habitat (*i.e*. forest vs open fields) affected the success ratio and prey attacks. The GPS tagged bats also predominantly caught prey in separate foraging bouts that were each dedicated to either hawking (Fig. 2A, blue diamonds) or gleaning (Fig. 2A, green circles). However, almost half of all aerial prey were captured during commuting (47 % of total aerial captures; Fig. S2). When gleaning, the bats attacked the same total number of prey in forest and open field habitats (field: 207 vs forest: 221 attack in total, Fig. 2C), but with more attacks per bout when gleaning above fields (field: 25 vs forest: 14 attacks/bout). Moreover, gleaning in open fields was twice as successful than in forest habitats (success ratio of 48 % per foraging bout with more than 2 prey attacks in open fields vs 12 % in the forest, Fig. 2E, Table S6). In contrast, when capturing insects in air, the attack rates and success ratios were consistently high and unaffected by habitat (Fig. 2DF, LMM, Table S5).

**Fig. 2:**
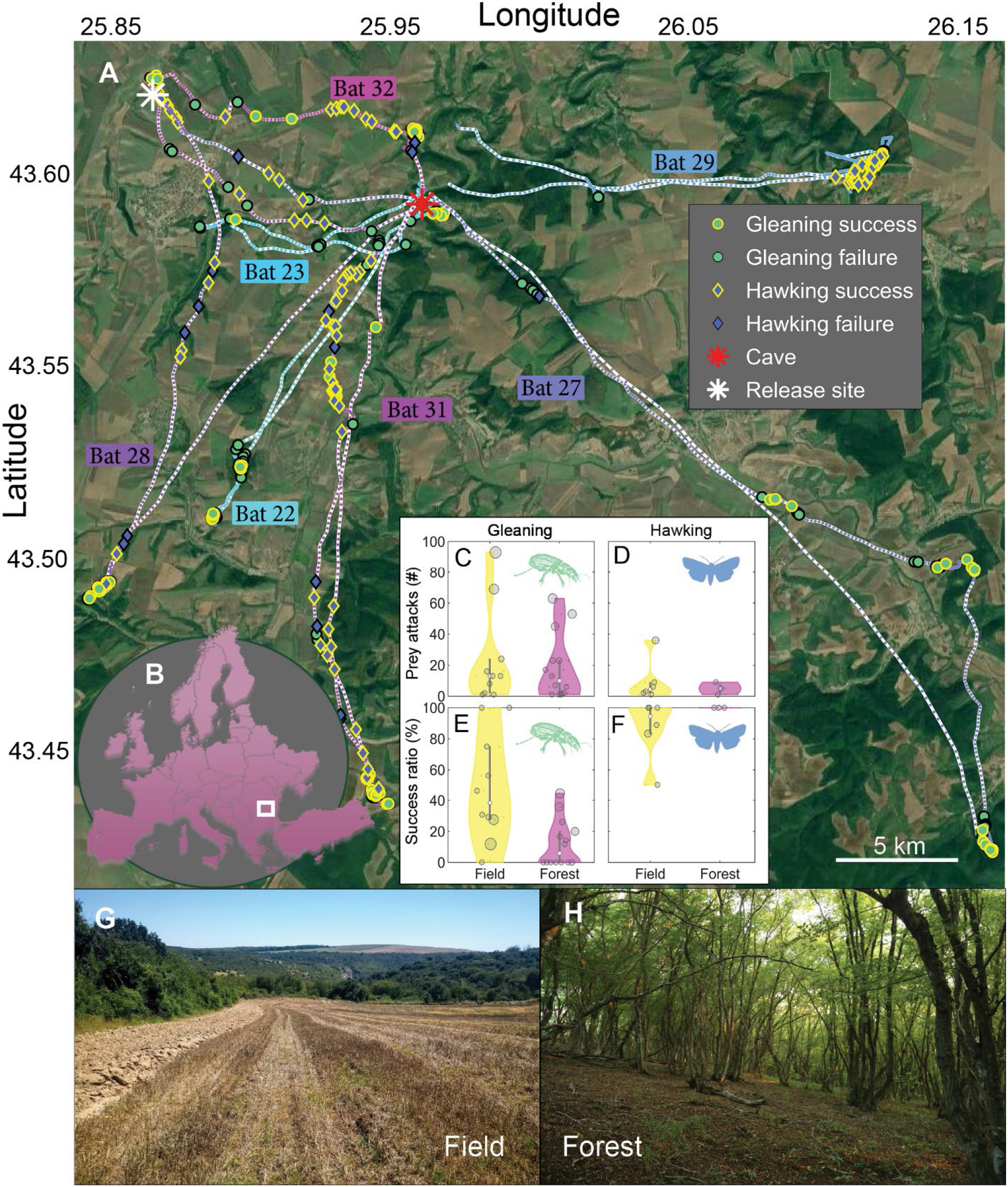
Habitat influences foraging success of greater mouse-eared only when gleaning. A) Tracks of seven bats with GPS tags released either at the cave (red star) or at a location nearby (white star) and their foraging behaviour: Gleaning (green circles) and hawking (blue diamonds) attacks with success (yellow edge) or failure (black edge). B) The bats were tracked in North-Eastern Bulgaria (white square). C-F: Total prey attacks (CD) and success ratios per foraging bout (EF), for both habitats: open field (yellow; G) and forest (magenta; H). Each data point corresponds to one foraging bout and is sized according to the number of attacks in the bout. G-H: The two main foraging habitats of greater mouse-eared bats: open fields (G) and the open spaces below the canopy in forests (H).

To estimate prey types and sizes, we first performed DNA metabarcoding analysis on the faeces of 54 untagged greater mouse-eared bats from the same colony caught in the morning upon returning from the foraging grounds. The bats target a wide range of prey species spanning 155 OUT (Operational Taxonomic Units) (Fig. 3AB), of which ~60 % occupy aerial niches (36 families) and ~40 % occupy ground niches (23 families; Fig. 3A). We measured the lengths of representatives of each species from online photo databases covering the same region in Bulgaria. Ground prey were 2.5x longer than aerial prey (20 mm (quartiles: 14–28 mm) vs 7 mm (quartiles 4–9 mm), Fig. 3C). We used length-weight regressions(Straus and Avilés, 2018) for each group to convert body-length to body mass for each prey type. For this analysis, we used the weighted average of the two most numerous prey types for aerial and ground regression values. Under these assumptions, estimated dry body masses of ground prey were ~20 times heavier than aerial prey (means: 67.5 mg (quartiles: 29.7–146.6 mg) vs 3.0 mg (quartiles: 0.7–5.7 mg, Fig. 3E, green and blue circles).

**Fig. 3:**
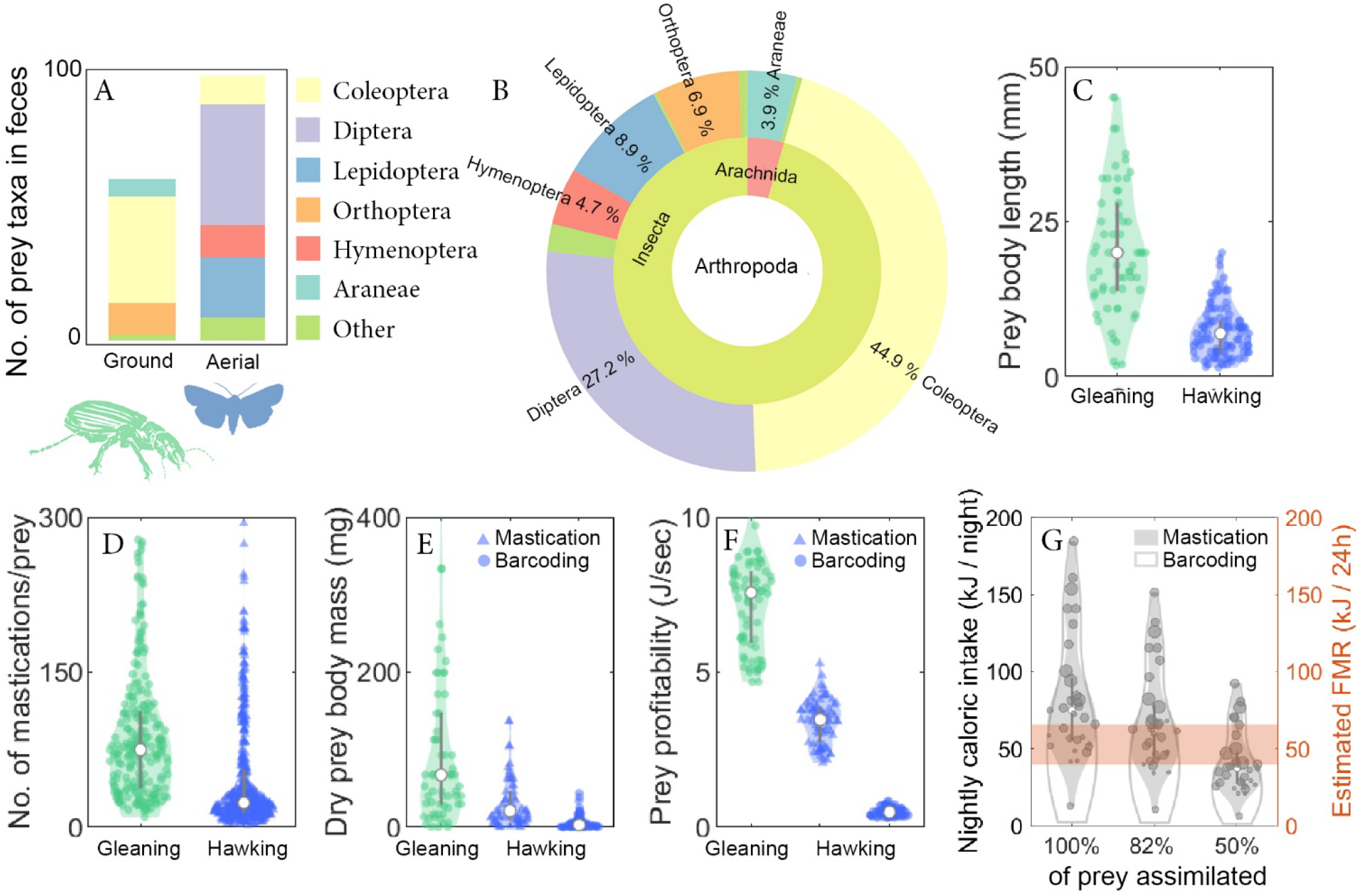
Ground prey is larger than aerial prey and sufficient to offset the lower foraging success ratios of gleaning. A-B: DNA metabarcoding of faeces from 54 greater mouse-eared bats (48 females, 6 males). Insects were categorized as either ground (green) or aerial (blue). The few species (N = 5) that are both aerial and ground were omitted from the analysis. Distribution of the targeted prey orders depicted as OTU (Operational Taxonomic Units) between ground (~40%) and aerial (~60%) niches (A) and across taxonomical units in the Arthropoda (B). C-F): Prey properties and profitability during gleaning (ground prey, green) and aerial hawking (aerial prey, blue), with kernel densities and boxplots. C) Body lengths of the prey sorted by foraging style. D) Number of mastication sounds identified after each prey capture by an automatic detector (N = 244 ground captures and 336 aerial captures across 10 bats). E) Dry prey body masses of each prey type identified for gleaning via DNA metabarcoding (green circles) (DNA metabarcoding was used as the reference prey body mass for ground captures), and for aerial prey by mastication analysis (blue triangles) and DNA metabarcoding (blue circles). F) Prey profitability of gleaning or hawking prey based on prey body masses from mastication analysis (triangles) or DNA metabarcoding (circles) combined with observed success ratios, handling and search times. The data are plotted for bootstrapped data (N=70 random data points) due to varying sample sizes of each parameter. G) Total caloric intake per night per bat calculated by multiplying the caloric intake per prey with the number of successful gleaning and hawking prey captures (Fig. 1), and compared to the field metabolic rate of a 30 g bat estimated from the literature (orange).

Since DNA metabarcoding does not provide the exact proportion of caught prey items and species, and thus does not allow to calculate the size distribution of caught prey, we performed an analysis of the mastication sounds as an additional proxy for prey size. Greater mouse ear bats chew all prey while flying irrespective of how they are caught and take longer to masticate larger prey (verified in laboratory feeding experiments, Fig. S9). We used body length to body mass conversions from the DNA metabarcoding of ground prey as the reference prey body mass. We then estimated aerial prey body masses from the difference in chewing durations between ground and aerial prey. Bats chew longer on ground prey than aerial prey, indicated by the ~3x more mastication sounds detected after each gleaning capture (75, quartiles: 38.5-111.5, Fig. 3D) compared to aerial hawking (23, quartiles: 15- 55). By applying this ratio to the body mass estimations of gleaning prey, we estimated aerial prey body mass of 21.2 mg on average (quartiles: 9.3-45.9) (Fig. 3E, blue triangles). Thus, in the following, we use both a lower and a higher estimate of aerial prey body masses of 3.0 mg (from DNA metabarcoding), and 21.2 mg (from masticating sounds) (Fig. 3E). Taking successful prey captures into account and the weighted average of the caloric values of the two most numerous prey types for aerial and ground (25.4 kJ/g dry mass of ground prey and 21.3 kJ/g dry mass of aerial prey), the bats ingested an average of 60.2 kJ/night/bat (quartiles: 32.1-84.8) based on the lower aerial prey body mass estimates (Fig. 3G, solid grey line), and 74.9 kJ/night/bat (quartiles: 55.2-95.7) based on the higher estimates (Fig. 3G, shaded grey). Using energy assimilation rates of 50-82% in bats(Kurta *et al*., 1989; Straus and Avilés, 2018), the bats obtained on average between 30-61 kJ/night per bat (Fig. 3G).

The profitability of prey caught by gleaning or hawking for all bats (N = 34) was quantified by combining success ratios (Fig. 1F), search and handling times (Fig. S4A) with lower (Fig. 3E, circles) and higher estimates of prey body masses (Fig. 3E, triangles). The gleaning foraging strategy yielded a profitability of 7.4 J/sec (quartiles: 5.6-8.1 J/sec, Fig. 3F green), while hawking resulted in a lower estimate of 0.5 J/sec (quartiles: 0.4-0.54, Fig. 3F, blue circles) and a higher estimate of 3.3 J/sec (quartiles: 2.8-3.8, Fig. 3F, blue triangles). Prey profitability when gleaning is thus 2.3-14 times higher than when aerial hawking.

## Discussion

The small size and high metabolic rate of bats, coupled with a costly locomotion mode, require an elevated and constant input of calories from foraging(Kleiber, 1947). This, in turn, calls for either stable and narrow food niches or adaptive hunting behaviours that track habitat dynamics. Here, we used biologging and metabarcoding to explore how greater mouse-eared bats chose between two different foraging strategies to cover their energy intake, and how strategy switching is adapted to habitat.

### Greater mouse-eared bats are more successful when hawking despite being gleaning specialists

Since gleaning requires passive listening, while aerial hunting requires vocalising(Arlettaz, 1996), we first hypothesized that the two strategies were mutually exclusive and would yield different prey attack rates and success ratios. Indeed, our data show that the tagged bats used both foraging strategies to capture food, but that foraging bouts were dedicated exclusively to either gleaning or aerial hawking (Fig. 1 & Fig. S11). Nonetheless, averaging over ten nights and individuals, tagged bats caught a mean of 25 insects in air and 30 insects on the ground during a night of foraging (Fig. 1), indicating a reliance on both food sources.

These estimated feeding intakes are slightly higher than the total prey captures of the similar-sized *Rhinopoma microphyllum*, estimated from buzz counts (Cvikel *et al*., 2015), but well below previous indirect estimates of feeding intakes in a smaller (7-11g) species (*M. daubentonii*) that were extrapolated to suggests that they should capture thousands of tiny insects per night. The discrepancy between the total prey captures could potentially be a consequence of the tag, or the tagging process. However, the extra weight of tags (~3-4 g)(Portugal and White, 2018; Kline, Ripperger and Carter, 2021) did not appear to strongly impact the ability of bats to capture food since i) both tagged and un-tagged trained bats quickly learned to intercept aerial and ground prey in the lab with similar success ratios as in the wild (Table S2, Video S1-2), and ii) the wild tagged bats spent the same amount of time on foraging outside the colony as bats equipped with lighter (0.4) telemetry transmitters (Egert-Berg, Hurme, Greif, Wilkinson, *et al*., 2018).

Given our measured total prey captures and prey sizes, wild greater mouse eared bats in our study assimilated an average of 61.4 kJ/night per bat (82 % assimilation). This is higher than the estimated field metabolic rate (FMR) of the similar-sized female lesser long-nosed bats (*Leptonycteris yerbabuenae*) of 40 kJ/day, but close to allometric scaling of the FMR of a 30 g bat based on heart rate measurements from wild-tagged 18 g *Uroderma bilobatum* (FMR_M.myo_ = FMR_U.bil_ * (30g/18g)^0.7^ = 65.5 kJ/day). Thus, despite making less than 100 prey captures per night, the estimated food intake of the tagged bats matches their predicted FMR. The discrepancy between the predicted intake of thousands of insects per night from a smaller bat species, and the measured total prey captures in our study is therefore more likely to relate to the dramatic difference in prey sizes between the prey species rather than a reduced foraging effort due to tagging effects. The bats in our study on average reached their predicted energetic requirement in a full night of foraging, but only just so, indicating that they may have little scope to compensate for changes in their environment. Since the bats fly out just after sunset and return early in the morning, they are vulnerable to any disturbance or change in habitat quality that reduces their foraging intake. Moreover, despite selecting only heavy, post-lactating females within the same colony, there was wide individual variation in hunting. Tagged bats attacked from 48 to 280 prey during a night of foraging (Fig. 1–2), demonstrating that continuous recordings from the same individuals are important to quantify the energy budgets of wild animals.

Even though the tagged bats captured the same number of prey by aerial hunting as by gleaning, we expected that their sensorimotor adaptation to gleaning would come at the cost of a poorer ability to capture aerial prey during hawking. Counter to our hypothesis, we found that hawking bats were highly successful (80%) compared to when gleaning (30%) (Fig. 1F). Despite that greater mouse-eared bats are gleaning specialists, we find that they have success ratios when hawking that are on par with observations in the wild for hawking bats (Rydell, McNeill and Eklöf, 2002). These high success ratios may be facilitated by superfast sensorimotor responses to guide echo-based capture. Such superfast movements may not benefit a gleaning strategy to the same extent since carabid beetles can seek refuge under leaves or twigs if the attack of the bat is not perfectly aimed on the prey. In such cases, the bats must rely on tactile and olfactory cues, and their poorer ground locomotion to find the prey(Kolb, 1958), which may explain the low success rate of gleaning (Table S2).

### Greater mouse-eared bats prefer larger ground-dwelling prey over aerial prey despite low success ratios

Tagged bats attacked prey in the air and on the ground at similar rates, but success ratios for aerial prey were more than twice of those of ground prey. Moreover, gleaning insects on the ground most likely exposes bats to a higher predation risk from ground predators, and a higher risk of injury (Toshkova et al. *In prep*; Brandmayr *et al*., 2009). This begs the question of why most of the tagged bats (N = 22 of 34 bats) nonetheless preferred to capture insects on the ground? To answer this we hypothesized that bats would choose the foraging strategy with the highest profitability (*i.e*. energy intake/time) (Stephens and Krebs, 1986). We estimated prey profitability by dividing estimated prey caloric values with prey search and handling time, factoring in the success ratios for each strategy. We find that gleaning prey off the ground, despite the much higher failure ratios, offers a prey profitability between 2.5 and 14 times higher than aerial hawking owing to the much larger ground prey (Fig. 3F). Beetles gleaned off the ground are 3-20 times heavier than aerial prey and have higher protein and fat content(Razeng and Watson, 2015). In fact 85 % of the energy returns from a night of foraging comes from gleaning despite the similar average number of prey capture attempts in air and on the ground per night. Thus the paradox of preferring low success ratios and higher risk of injury while gleaning (Fig. S10) is explained by a much higher prey profitability: the bats opt for a high risk-high gain strategy. Nevertheless, we find that aerial hunting remains as a valuable supportive foraging strategy for 19 out of 34 individuals on the nights sampled (Fig. 1DE), contributing 15 % of the total intake for these bats. This reliance on aerial hawking next prompted us to investigate how habitat affected prey profitability and the choice of foraging strategy.

### Prey profitability changes with habitat, but does not affect foraging decisions in greater mouse-eared bats

It has been hypothesized that greater mouse-eared bats are opportunistic predators able to maximize their energy intake by choosing the most profitable habitat (Arlettaz, 1999). We tested this hypothesis by investigating how prey habitat (*i.e*. open fields vs forests (Arlettaz, 1999), Fig. 2GH) or movement style (i.e. commuting or active searching for prey, Fig. S2) affect the profitability of prey and thereby influence foraging decisions of the wild bats (N = 7 GPS-tagged bats). The bats primarily gleaned prey during dedicated foraging bouts (Fig. S2A), whereas hawking was used equally during commute and in foraging bouts (Fig. S2B). This indicates that aerial hawking is a flexible strategy that is efficient enough to use during commuting. This perhaps offsets some of the energetic costs of the often lengthy commutes made by greater mouse-eared bats to preferred foraging areas while also potentially providing them with concurrent information about aerial prey profitability.

When actively gleaning for prey, the tagged bats were more successful and attacked more prey per bout in open fields than in forest habitats(26 vs 15 attacks/bout), but performed more foraging bouts in the forest (field: 8 foraging bouts vs forest: 15 bouts; 3 bouts were a mixture between field and forest). Thus, the quality of the gleaning habitat (here defined as a habitat enabling higher success ratios and attack rates for the same strategy) does not appear to affect the decision of which habitat to target. This could have several explanations: bats may be maximizing energy intake by switching to forest habitat after depletion of open field habitats, or they could be balancing predation risk and/or conspecific competition which are both likely to be higher in open fields. In contrast to gleaning, bats were equally successful when using echolocation to capture insects above open fields and below the canopy in forests revealing that these bats are well able to hunt in semi-cluttered spaces(Stidsholt *et al*., 2021). Such a flexible strategy allows them to be efficient hunters across different habitats equipping them to exploit diverse environments.

Although this bat species hunt independently, the dominant foraging strategies of bats tagged on the same night were more similar than for bats tagged on different nights. This indicates that temporally-varying environmental parameters such as rain, wind or mass emergence of aerial insects, experienced by all bats in the area on a given night, influence the most beneficial foraging strategy. For example, after rain, rustling sounds of walking arthropods on leaves are more difficult for bats to detect potentially halving the detection range(Goerlitz, Greif and Siemers, 2008). Reduced detection distance could not only lower detection rates but also capture success ratios (*i.e*. less time to plan and execute a capture). The time between gleaning prey attacks increased by an average of 35 seconds on the two nights with the most aerial captures (22^nd^ of July 2018 and 2019, GLMM; testing whether the time between gleaning attacks was explained by nights with a majority of aerial captures, p<0.01) and the bats switched more often between gleaning and aerial strategies on an attack-to-attack basis (GLMM; testing if the number of switches per night was explained by the nights with a majority of aerial captures, p<0.01). Longer periods between prey attacks and more strategy switching indicate that gleaning prey on these nights was less profitable compared to aerial prey, either due to poor conditions on the ground or a mass emergence of aerial prey.

Our results demonstrate that greater mouse eared bats often resort to aerial hawking despite their preference and specialisation for ground gleaning. Although much smaller prey are taken on-the-wing, the consistently high success ratios and prey captures rates of hawking independent of habitat make this a reliable backup strategy. The same qualities may make hawking a more widespread strategy than expected in other gleaning bat species. Such foraging flexibility both adds to the energy intake of wild bats and also improves their resilience to changing conditions. Thus, this strategy might have allowed a wide range of bat species to tap into the unpredictable, ephemeral food resources in open spaces. This would have put an evolutionary pressure on maintaining aerial hawking capabilities to secure an additional food resource in fluctuating environments, and may also explain why hawking remains as a foraging strategy in many bat species traditionally seen as gleaning and trawling specialists.

## Conclusion

We show that greater mouse-eared bats, a gleaning specialist, catch relatively few prey items per night on the ground at high failure ratios, but still achieve a high prey profitability by targeting large, energy-rich prey. This shows that prey availability and size, weighted by relative hunting success, are important drivers of foraging decisions in wild bats. We find that the bats use gleaning as a primary foraging tactic in a high risk-high gain approach, and switch to aerial hunting when environmental changes reduce the profitability of ground prey. We conclude that prey switching matched to environmental dynamics plays a key role in covering the energy intake even in a specialised predator.

## Material and Methods

### Method details

All experiments were carried out under the licenses: 721/12.06.2017, 180/07.08.2018 and 795/17.05.2019 from MOEW-Sofia and RIOSV-Ruse. We tagged and recaptured 34 female, post-lactating greater mouse-eared bats with sound-and-movement tags from late July to mid-August in the seasons 2017, 2018 and 2019. The bats were caught with a harp trap at Orlova Chuka cavfe, close to Ruse, NE-Bulgaria, in the early mornings as they returned to the roost. The bats were kept at the Siemer’s Bat Research Station in Tabachka to measure the forearm lengths, CM3 and body weights (Table S1). Bats weighing above 29 grams were tagged and released the following night between 10-11 p.m. at a field 8 km from the roost (Decimal degrees: 43.622097, 25.864917) or in the colony. The tags were wrapped in balloons for protection and glued to the fur on the back between the shoulders with skin bond latex glue (Ostobond). The bats on average spend 2 to 14 days equipped with the tags until we recaptured the bats at the cave or the tags detached from the bats and fell to the ground below the colony. Upon recapture, the bats were weighed and checked for any sign of discomfort from the tagging before they were released back to the colony.

### Tags

We used two different tags for this study. Both tags recorded continuous data during one night of foraging. The first tag (Tag A) recorded audio data with an ultrasonic Knowles microphone (FG-3329) at a sampling rate of 187.5 kHz, with 16 bit resolution, a 10 kHz 1-pole analog high-pass filter and a clipping level of 121 dB re 20μPa pk. These tags also included triaxial accelerometers sampling the movement of the bats at 1000 Hz with a clipping level of 8 g. All accelerometer data were calibrated, converted into acceleration units (m/s^2^) and decimated to 100 Hz. The orientation of the bats were recorded with triaxial magnetometers using a sampling rate of 50 Hz. These tags weighed from 3.5-3.9 g (including VHF for localisation and recapture of the bats). The second tag (Tag B) recorded audio using a MEMS microphone sampling at 94 kHz and with 16 bit resolution. The movement of the bats were recorded with triaxial accelerometers at 50 Hz sampling rate and with a clipping level of 8 g. These tags also included GPS sensors that logged the position of the bats every 15 seconds. These tags weighed 3.9-4.2 g (including VHF). In total, we have data from 16 bats with Tag A and 18 bats with Tag B (of which 5 are 50 % duty cycled) (Table S1).

### Tagging effects

Both tags weighed 11-15 % of the bodyweight of the tagged bats. We addressed the effects of tagging on the data and the bats by the following procedures: (i) We trained bats in a flight room to capture mealworms and moths tethered on strings and glean beetles of different sizes from either a bowl or a square meter of natural forest floor (Table S2). The bats caught aerial prey with high success ratios (95 % for mealworms and 69 % for moths (N = 2 bats)) with tags and without tags (75 % for mealworms (N = 4 bats)). The bats gleaned beetles with a success rate of 39 % from the square of forest floor. The success rate increased to 85 % when they were catching beetles from a bowl with no escape options for the prey (N = 3 bats). This indicates that the low gleaning success ratios in the wild are most likely not caused by tagging effects, but more likely because beetles can escape in cracks or below leaves. Additionally, we could not detect any visual difficulties with capturing prey from strings or beetles from the ground (Video S1-2). (ii) The wild bats caught prey with high success ratios in air indicating that the tag effects had little or no impact on the ability to capture prey. (iii) A previous study with the same species and tags found that tagged and untagged bats spend equal amount of time foraging (Egert-Berg, Hurme, Greif, Wilkinson, *et al*., 2018). (iv) The weight loss of these bats of 3-4 % during the instrumentation time was equal to the weight loss of control bats carrying only VHF radio transmitters (0.5 g) indicating that handling and carrying of tags might disturb the bats, but the additional extra load did not seem to add further energetic consequences to the bats in addition to the VHF (Egert-Berg, Hurme, Greif, Goldstein, *et al*., 2018).

## Quantification and statistical analysis

To understand how the tagged bats allocated their time and captured prey in the wild, all wild tag recordings were manually analysed by displaying the acoustic and the movement data in 7-20 second segments with an additional option of playing back audio data. The visualisation included three separate windows with synchronized data: (i) An envelope of audio data filtered by a 20 kHz 4 pole high pass filter to detect the echolocation calls. (ii) A spectrogram of audio data filtered by a 1 kHz 1 pole high pass filter to visualise the full bandwidth acoustic scene showing echolocation calls, conspecific calls, chewing sounds, wind noise etc. (iii) The final window showed triaxial accelerometer aiding the identification of wingbeats, landing, take-offs as well as capture events.

### Categorisation of capture attempts

Greater mouse-eared bats are known to glean prey off the ground and to capture aerial prey. To recognize gleaning capture attempts in the wild data, the tag recordings were ground truthed by analysing sound and movement data from capture attempts of bats under controlled experimental settings in the lab. Two individuals were trained to catch walking beetles on vegetation using passive hearing while carrying a tag (Video S1). These ground captures were identified by stereotyped patterns consisting of three simultaneous events (i) Low vocalizations (around 50 dB re 20μPa^2^s) prior to the capture attempt indicating that the bat was using passive listening to listen for prey generated cues. (ii) A short, broadband and loud audio transient simultaneous to a peak in the accelerometer data indicating that the bat was landing on the ground. (iii) The accelerometer signal indicating a landing was an increase in wingbeat frequency and amplitude prior to the landing, a peak in the sway and heave axis (y and z dimension, often with opposite values) at the time of contact with the ground and often followed by a flattening of the signal on all three axes. These stereotyped audio and accelerometer signals found in the laboratory experiments were matching signals seen in the wild data (Figure S5). These signals were then identified in all wild tag recordings during the visualisation and marking process. Aerial capture attempts were identified in the wild data if a buzz was present. Only buzzes in flight were marked to exclude landing buzzes. In addition, each capture attempt was marked as “successful” or “unsuccessful” based on the presence or absence of chewing sounds. The chewing sounds were audible (Sound files are uploaded) and for the low-noise tag recordings visible in spectrograms (Figure 3). For the five 50 % duty cycled tag recordings we doubled the foraging attempts and successes.

### Behavioural analysis

To evaluate time allocation, we analysed data from 15 tags with both accelerometer and magnetometer data. We separated times of rest from flight through identification of wingbeat epochs. Wingbeats were detected as cyclic oscillations in the z-axis dimension of the accelerometer data. We first band-pass filtered the z-axis dimension of accelerometer data (from 5 to 25 Hz) by a delay-free symmetric FIR-filter (filter length: 1024 samples, sample rate: 100 Hz). We then identified flight epochs as the time intervals where a running mean of 50 seconds of the wingbeat data were above a threshold of 20 m/s^2^. A window length of 50 seconds was chosen to avoid short, flight epochs consisting of only few wingbeats.

We identified times of foraging from travelling during flight epochs by changes in heading because the bats fly straight towards and between foraging grounds (Egert-Berg, Hurme, Greif, Goldstein, *et al*., 2018). Heading was computed by gimballing low-pass (3 Hz) filtered triaxial magnetic field measurements with the pitch and roll estimated from down-sampled and low pass filtered (3 Hz) accelerometer data (Johnson and Tyack, 2003). We applied a running mean of 50 seconds to the heading measurements to evaluate foraging bouts. We chose a length of 50 seconds similar to the minimum foraging bout length used by Hurme et al., (2019) and in our GPS analysis. Foraging bouts were identified as time intervals where the envelope of the signal was above a threshold of 0.05. Due to the large variation between animals, this threshold was raised to 0.3 for six tag recordings to avoid more than 10 switches between travelling and foraging per night as used by Hurme et al., (2019) and in our GPS analysis. We omitted identified foraging bouts with no capture attempts from the analysis (N = 62 out of 202 foraging bouts for 16 bats).

### GPS and habitat analysis

We recaptured 7 tags with GPS data. We used first-passage time analysis (Fauchald and Tveraa, 2003) to identify foraging bouts from travelling bouts (Hurme *et al*., 2019) by using the R package “adeHabitatLT” (Calenge, 2006). First, we converted the latitude and longitude coordinates from degrees to meters (“proj4string” function in adeHabitatLT) and then regularised the GPS tracks to 10 second time stamps. We then calculated the first-passage times for all radii ranging from 5 to 400 m (in 5 m steps). We plotted the variance of the log-transformed first-passage times and found the highest value of around 250 m for all bats which is in accordance with a previous study on *Myotis vivesi* (Hurme *et al*., 2019). This value estimates the scale at which the bat is operating, and was used for all tracks. We used the Lavielle method (“lavielle” in adeHabitatLT) to divide the path segments into foraging and non-foraging bouts (Calenge, 2006). The minimum number of locations in a bout was chosen to 5 (corresponding to 50 seconds), and the maximum number of segments per night was 50. The function “chooseseg” was chosen to find the number of segments at which the contrast between bouts were highest. The number of segments per night was estimated to either 10 or 11 per bat. To find a threshold value to separate foraging and travelling bouts, we found that the distance travelled per segment showed bimodal distribution (function “bimodalitycoeff” in Matlab (Zhivomirov, 2022)). We then fitted a Gaussian mixture model with two components to the data (Figure S6). We defined the threshold between the two distributions as the lowest quartile of the non-foraging (*i.e*. travelling) segments (Figure S6) corresponding to 40 meters travelled per 10 second segment.

To determine in which habitats the bats were foraging, we transferred all tracks onto a Google Map Satellite Imagine (using: “plot_google_map” in Matlab version 2021b). We manually determined whether the GPS locations for each segment were located in one of three categories: Field, Forest or both/others. We excluded foraging (N = 4) and non-foraging (N=23) bouts that were not assigned to either field or forest.

### Statistical analysis

All statistical modelling was performed in R (version 4.0.3). We fitted different models to understand how capture attempts and success ratios were influenced by the foraging strategy, the habitat and the mode of action (commuting vs foraging in bouts). For all models, we used a goodness of fit evaluation based on the marginal (R_(m)_^2^) and the conditional R2 (R_(c)_^2^) (Nakagawa and Schielzeth, 2013), the 0.05 criterion for statistical significance and Bat ID as random effect. We examined potential collinearity between the predictor variables of each model using variance inflation factor (from R-package “car” (Fox and Weisberg, 2019)). No collinearity was found.

#### Model 1

We investigated how capture attempts and foraging successes were influenced by the foraging strategy (N=29, the five 50 % duty cycled tags were omitted from the analysis). We used foraging strategy as predictor variable and fitted two linear mixed effect model to the data (“lme4” R-package (Bates *et al*., 2015)). In the first model (1a), we used foraging attempts as response variable assuming a Poisson distribution (link=”log”), and in the second model (1b), we used foraging success ratios as a normally distributed response variable. We examined the residuals (“DHARMa” package in R) and found no deviations from the expected distribution. Model 1a showed that the amount of capture attempts in a night was explained mostly by the individual bat (random effect explained 70 % of the variance of the data) and less by the foraging strategy (27 %) (Table S3). Overall, model 1b explained 77 % of the deviance of foraging success. Subtracting the random effect of individual bat only decreased the explained deviance by 8 % (Table S4). Model 1b revealed that bats are more successful when hawking for prey.

#### Model 2

We investigated how the capture attempts and foraging successes were influenced by the habitats of the seven bats tagged with GPS tags (Tag B). For both models, we used three categorical predictor variables: habitat type (field vs forest), foraging strategy (aerial vs hawking) and movement style (commuting vs foraging in bouts).

In model 2a, we used capture attempts (sum of failed and successful capture attempts in each foraging bout) as a response variable and fitted a GLMM (“glmer” function in R-package “lme4”) to the data with a Poisson distribution of the response variable and a link “sqrt” function. In model 2b, we used foraging success (per foraging bout) as a normally distributed response variable and fitted a LMM to the data (“lme” function in R-package “nlme” (Pinheiro *et al*., 2022)). We used model selection procedures (“dredge” in R-package (Burnham and Anderson, 2002)) to examine the best-fitted models using the AICc (corrected Akaike information criterion). The best-fitted models included all three predictor variables in both model 2a and 2b. We examined the residuals (“DHARMa” package in R) and slight deviations from the expected distributions were found.

In model 2a, the three predictor variables only explained 15.0 % of the data, while the random effects explained 78.1 % of the deviance in capture attempts. Thus, model 2a revealed that the variance in capture attempts per night is largely explained by individual bat rather than habitat, strategy or movement style (Table S5). In model 2b, the three predictor variables explained 72.3 % of the deviance of the success ratios. No difference was found when subtracting the random effect for model 2b. This model revealed that habitat and strategy strongly influence the success ratios of the bats (Table S6).

#### Model 3

We tested whether the immediate environment affected the foraging decision in bats by investigating the relationship between the dominant foraging strategy and the night of tagging for all 34 bats. The dominant foraging strategy was found as the ratio between aerial and foraging attempts ranging from 0 to 1 (0 meaning 100 % aerial foraging; 1 meaning 100 % gleaning) and was used as a normally-distributed response variable. The data was fitted with a LMM (function: “nlm” in R (Pinheiro *et al*., 2022)) using release night as a categorical fixed effect, and release site as a categorical random effect. This model showed that there was strong evidence of individual night on the chosen foraging strategy, indicating that bats on the same nights choose the same foraging strategy. (N = 10 different nights, 34 bats, and a mean of 3 (2.1 SD) tags per night (Figure S2, Table S7)).

### Relative prey sizes based on quantification of mastication sounds

After the visualisation and marking of all capture attempts, a custom-written chewing detector was used to automatically identify mastication sounds for nine tag recordings with sound quality to perform this analysis. This analysis included 452 ground captures attempts and 387 aerial capture attempts. The detector was used in two steps: i) To automatically determine whether the capture attempt was successful or not to verify the manual decision process based on listening to the chewing sounds. ii) To characterise the mastication sounds of each prey item from successful captures.

We extracted mastication sounds when the bat was flying after each capture attempt. Since the bats in flight chewed between echolocation calls, we analysed the intercall interval from the time of capture to either next capture attempt or 100 seconds ahead. We first detected mastication sounds and then classified the mastication sounds for each intercall interval.

Each intercall interval was extracted, filtered by a 7 to 15 kHz 4-pole Butterworth filter and convolved with a 40 ms Hanning window to exclude transients. These parameters were based on mastication sounds from capture bats with peak frequencies of 7 kHz which corresponds to the same peak frequency in wild bats. The intercall interval was classified as containing a mastication sound if the maximum amplitudes of the filtered signals were above a threshold of 0.012 and the peak frequency was above 5 kHz and below 20 kHz (Figure S7-8 black vs red). The intercall intervals that included mastication sounds were then used for the classification. Here we filtered the intercall intervals of the original sound data containing mastication with a 5 kHz 4-pole high-pass filter to reduce flow noise. To extract the onset of chewing as well as the duration, the detector also automatically extracted the time at which the bat produced the 10^th^ and 90^th^ quantile of the chewing sounds (Fig S7-8, grey dashed lines). The 10^th^ and 90^th^ quantile were a conservative choice to avoid false detections. The onset of the chewing was determined when the bat emitted the first sound in this interval. The duration of the chewing was determined as the length of this interval. The handling time was estimated as sum of the onset and duration of the mastication.

To test the performance of the classifier, the automatic and manual classification of successful vs unsuccessful prey captures were compared. A confusion matrix was made separately for all ground and aerial capture attempts. The classifier was evaluated by calculating the positive predictive value (PPV) and the false negative rate (FNR) based on the derivatives of the confusion matrix from the classification of the ground and aerial capture attempts (Table S8). Overall, the detector worked with PPV values above 0.99. However, the FNR was higher for aerial captures (3 % for ground; 9 % for aerial captures). The maximum sound energy in chewing after aerial captures were weaker than after ground captures which may explain the worse performance in air. To verify the mastication detector, we extracted and characterised mastication sounds of known prey types and sizes in controlled setups in the laboratory where the bats were eating while flying after prey captures (Figure S9).

### DNA metabarcoding

To understand the taxonomic diversity of the diet of wild mouse-eared bats we collected faecal samples from 54 bats (n= 26 in 2017, n= 28 in 2019, n = 48 female bats) returning to the roost after foraging. Individual bats were caught with a harp-trap positioned at the entrance of Orlova chucka cave, Pepelina, Bulgaria and placed in individual clean cotton bags until defecating. Faecal samples were collected in 98% alcohol and stored until further analysis. DNA extraction, data sequencing and bioinformatics were done following Morinière et al., 2016 (see also Morinière et al., 2019). In short, DNA from the faecal samples were extracted by using the DNEasy blood & tissue kit (Qiagen) following the manufacturer’s instructions. Multiplex PCR was performed using 5 μL of extracted genomic DNA and high-throughput sequencing (HTS)-adapted mini-barcode primers targeting the mitochondrial CO1 region. HTS was performed on an Illumina MiSeq v2 (Illumina Inc., San Diego, USA) at AIM - Advanced Identification Methods GmbH, Leipzig, Germany. Further, FASTQ files were combined and sequence processing was performed with the VSEARCH v2.4.3 suite (Rognes et al., 2016) and cutadapt v1.14 (Martin, 2011). Quality filtering was performed with the fastq_filter program of VSEARCH, fastq_maxee 2; a minimum length of 100 bp was allowed. Sequences were dereplicated with derep_fulllength, first at the sample level and then concatenated into one fasta file, which was then dereplicated. Chimeric sequences were then detected and filtered out from the resulting file. The remaining sequences were clustered into OTUs (Operational Taxonomic Units) at 97% identity. To reduce likely false positives, a cleaning step was employed that excluded read counts in the OTU table of less than 0.01% of the total read number. OTUs were blasted against a custom Animalia database downloaded from BOLD (Barcode Of Life Database, www.v3.boldsystems.org) and BIN (Barcode Index Number) information.

We measured the lengths of representatives of each species from online photo databases (www.boldsystems.org) or the reference collection of The National Museum of Natural History Sofia (Bulgaria) covering the same region in Bulgaria. The lengths used for the calculations were estimated as the maximum values of the measured prey lengths.

### Caloric value estimations

To estimate the energetic intake of one night per bat, we first used length-weight regressions to covert arthropod body-lengths to body masses (Straus and Avilés, 2018). Here we used the weighted average of the two primary arthropod orders from each foraging strategy: ground diet (*i.e*. Carabidae (78 %) and Orthoptera (22 %)) and aerial diet (*i.e*. Diptera (65 %) and Lepidoptera (35%)) (Straus and Avilés, 2018). From this calculation, we estimated the dry body masses of ground and aerial prey only based on metabarcoding data. In addition to this, we also calculated the dry body mass of aerial prey based on the ratio between the mastication (number of mastications/capture) after ground and aerial prey. This gave us an additional value for the dry body mass of aerial prey. Thus, we proceeded with a higher and lower estimate of aerial prey dry masses in the following calculations.

To convert dry body masses (mg/prey) into caloric values (J/prey), we multiplied the dry body masses of the prey with the caloric values for either ground (Bell, 1990; Zygmunt, Maryansky and Laskowski, 2006) or aerial (Kurta and Kunz, 1987; Bell, 1990) prey. Here, we used the caloric values of the weighted average of the two most numerous prey orders for ground and aerial prey. Nightly caloric intake of each bat was then calculated by multiplying the number of eaten ground and aerial prey items with the caloric values of each prey type.

### Profitability index calculations

To compare the profitability between the two foraging strategies, we first calculated the relative profitability of each prey without including prey size using the following relationship:

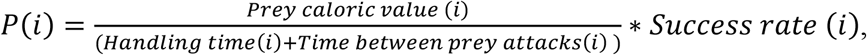

*i* = foraging strategy, Success rate (Mean success rate per bat (per bout with more than one capture) as a fraction from 0 to 1), Time between prey attacks (mean number of seconds between prey attacks per bat per night with unit seconds). Prey caloric values was estimated using both metabarcoding and mastication analysis (see *Caloric value estimations*). The prey profitability unit is therefore J/second per prey capture.

## Acknowledgements

We are thankful to Kaloyana Kosseva for help with bat feces collection in the field and to Ilias Foskolos and Nor Amira Abdul Rahman for assistance with laboratory experiments. We are grateful to the entire crew at the Siemers Bat Research Station for the support during the seasons 2017-2019 and to the Directorate of the Rusenski Lom Nature Park, Bulgaria.

## Author contributions

L.S. was responsible for tagging data collection and analysis, interpretation, and drafting of the manuscript. S.G. was responsible for conceptualization, tagging data collection, and interpretation of the data. A.H. collected the DNA metabarcoding data, analyzed and interpreted the data. H.R.G. was responsible for conceptualization and interpretation of the data. M.J. designed and manufactured the tags and contributed to the interpretation of the data. Y.Y. designed the tagging experiment and was responsible for conceptualization. P.T.M. was responsible for conceptualization and interpretation of the data. All authors contributed to the writing of the manuscript.

