## Supplemental methods for "Echolocating bats prefer a high risk-high gain foraging strategy to increase prey profitability"

### Extended data

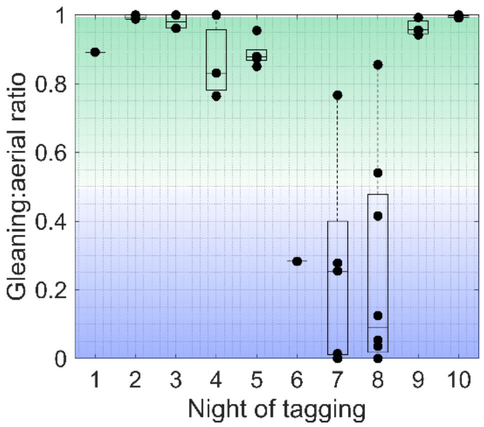


**Figure S1: The night of tagging affected the dominant foraging strategy of each bat.** For each night of tagging, the ratio between the number of gleaning and aerial prey capture attempts were calculated. A ratio of one indicates that the bat only encounters gleaning prey (solid green); a ratio of zero indicates purely aerial prey encounters (solid blue).

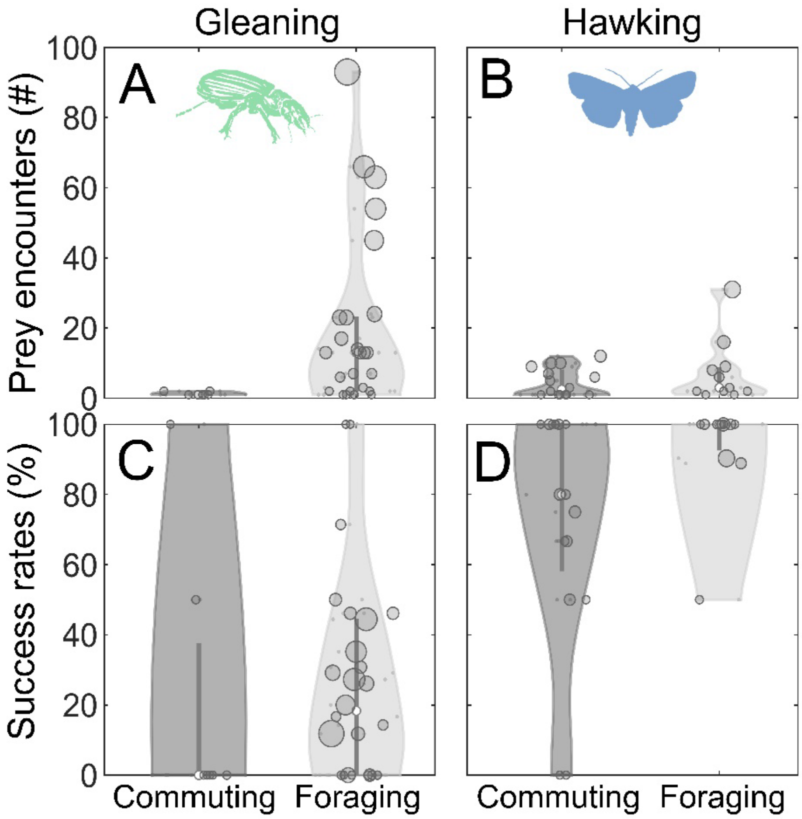


**Figure S2: Foraging success and prey encounter rates according to movement style.** The movement style was divided into either commuting to and from foraging sites and the cave (Commute), or foraging in foraging bouts (Foraging).

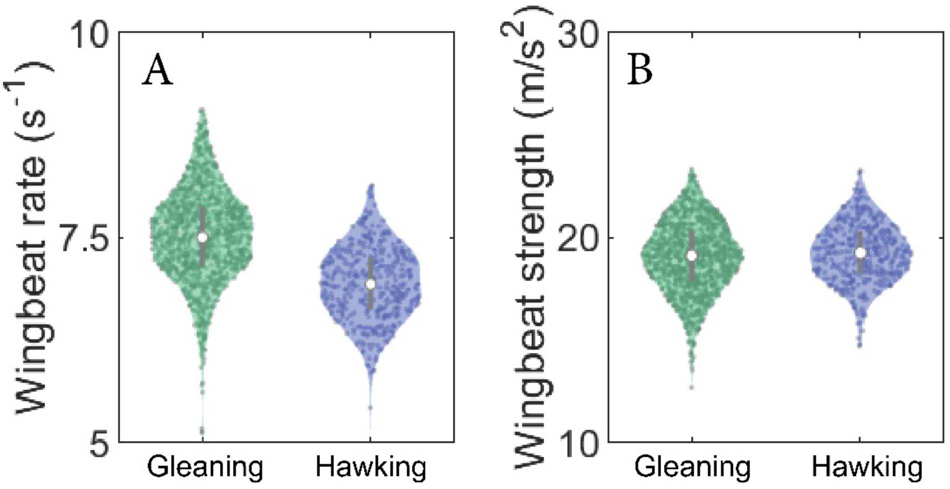

**Figure S3: Wingbeat rate and strength are similar across foraging strategies.** All data are plotted for aerial hawking (blue) and gleaning (green) along with kernel densities and boxplots. A) Mean wingbeat rate of the time interval between each prey encounter. B) Wingbeat strength calculated as the mean of the peak acceleration of each wingbeat in the time between prey encounters.

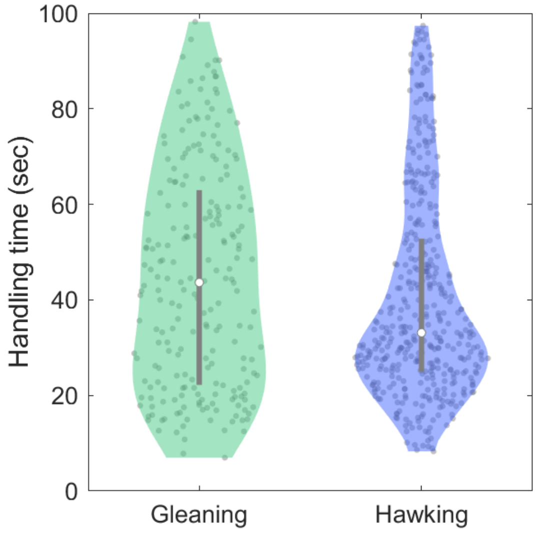


**Figure S4: Handling time of the prey caught either by gleaning or hawking is similar.** Handling time was estimated from the end of the prey capture to the end of the mastication.

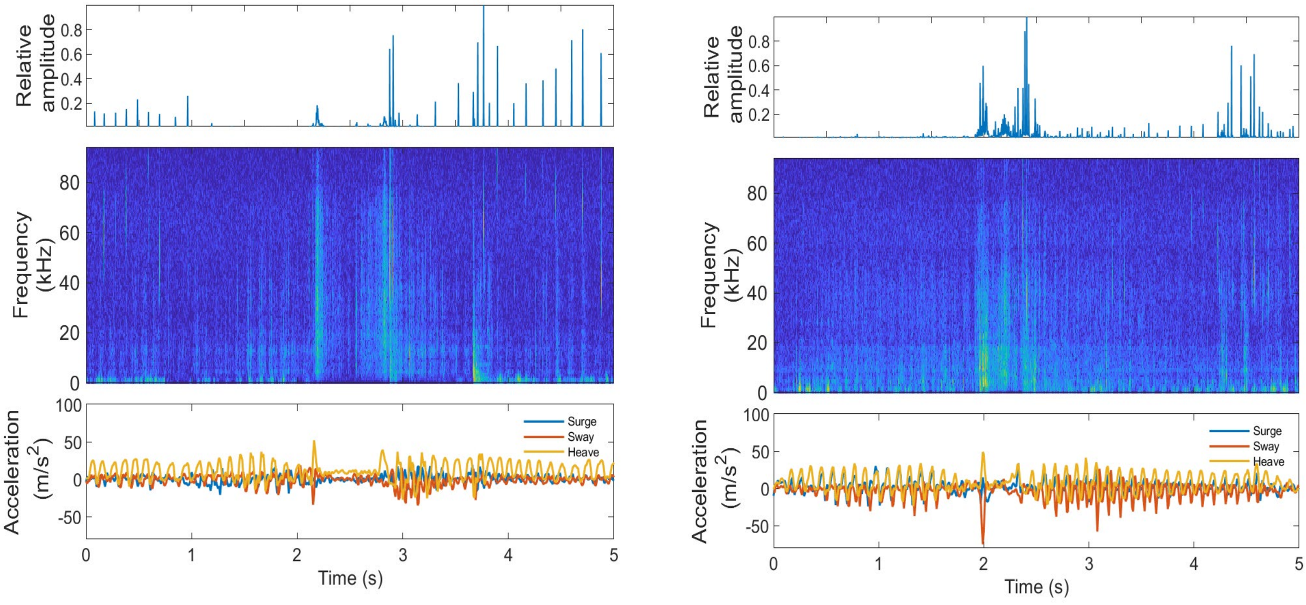

**Figure S5: Visualisation of ground capture attempts.** Synchronized audio and accelerometer signals during ground capture attempts are shown in the wild (left) and in a laboratory setup (right). The audio signals (upper two panels) are characterised by low output levels before the capture event. In contrast, the output levels increase afterwards. When the bat lands on the ground (at around time = 2 s in both figures), a loud and broadband acoustic “landing” signal appears. At the same time, a stereotypical acceleration pattern marks the landing, taxi and take-off (lower panel). The peaks in the heave and sway dimensions (yellow and red) mark the time at which the bat lands on the ground.

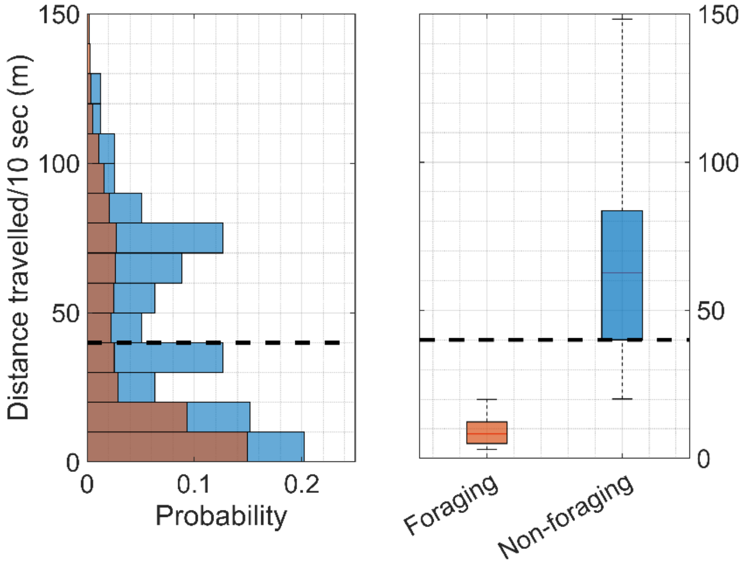


**Figure S6: Foraging vs non-foraging bouts distinguished based on the bats travelling below or above 40 m per 10 sec track.**

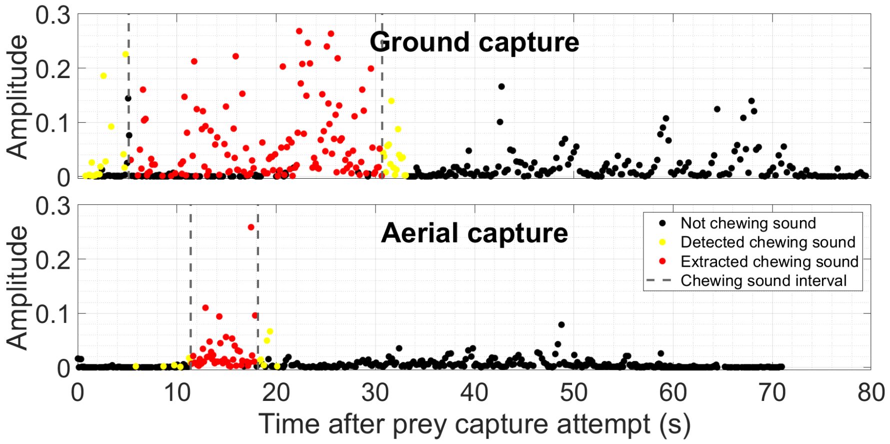


**Figure S7: Illustration of the automatic detector used for chewing sounds after successful prey capture attempts in the wild.** The intercall intervals of all calls emitted after a prey capture attempt and until next prey encounter have been extracted, band-pass filtered (5 to 15kHz) and convolved with a 40 ms Hanning window. The maximum of these sounds are plotted in black. If the maximum value was above the threshold of 0.012 and the un-processed intercall interval had a peak frequency within 5 to 20 kHz, these sound clips were identified as potential mastication (yellow). The 10$^{th}$ to 90$^{th}$ quantiles of the mastication sounds were extracted (grey dotted lines). Only detected chewing sounds within this interval were identified as mastication (red dots).

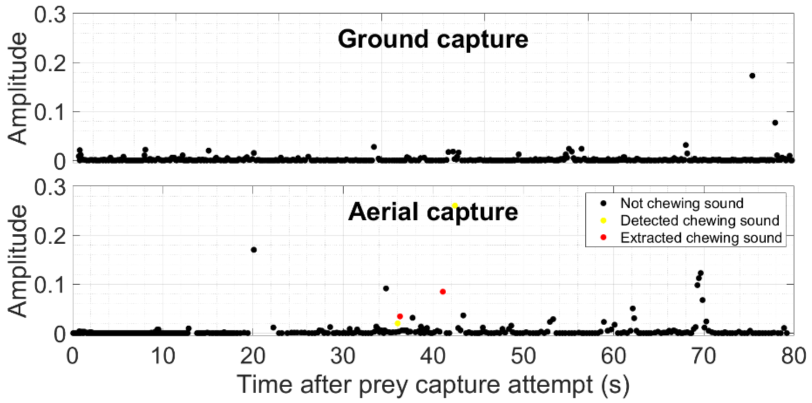


**Figure S8: Illustration of the automatic detector used for chewing sounds after unsuccessful prey capture attempts in the wild.** The number of extracted chewing sounds (red dots) are below four and thus classified as an unsuccessful prey encounter

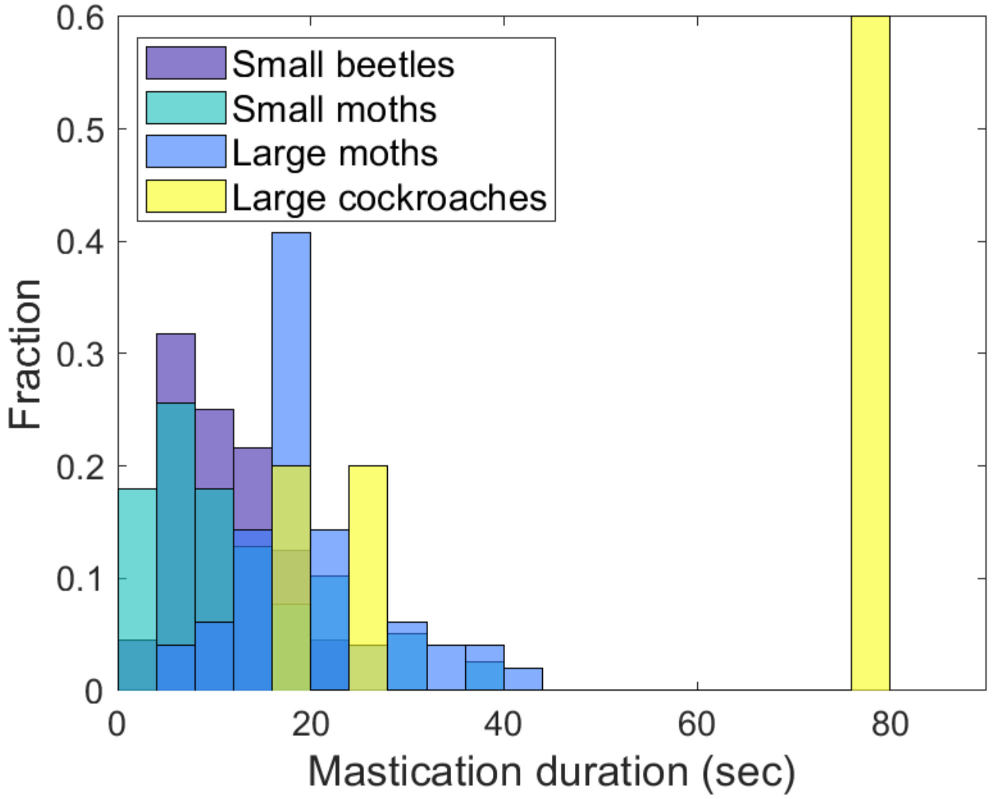


**Figure S9: Durations of chewing for different insect types and sizes.** The duration of the chewing was dependent on the size of the known prey types in the laboratory experiments: The bigger the prey, the longer the duration of the chewing.

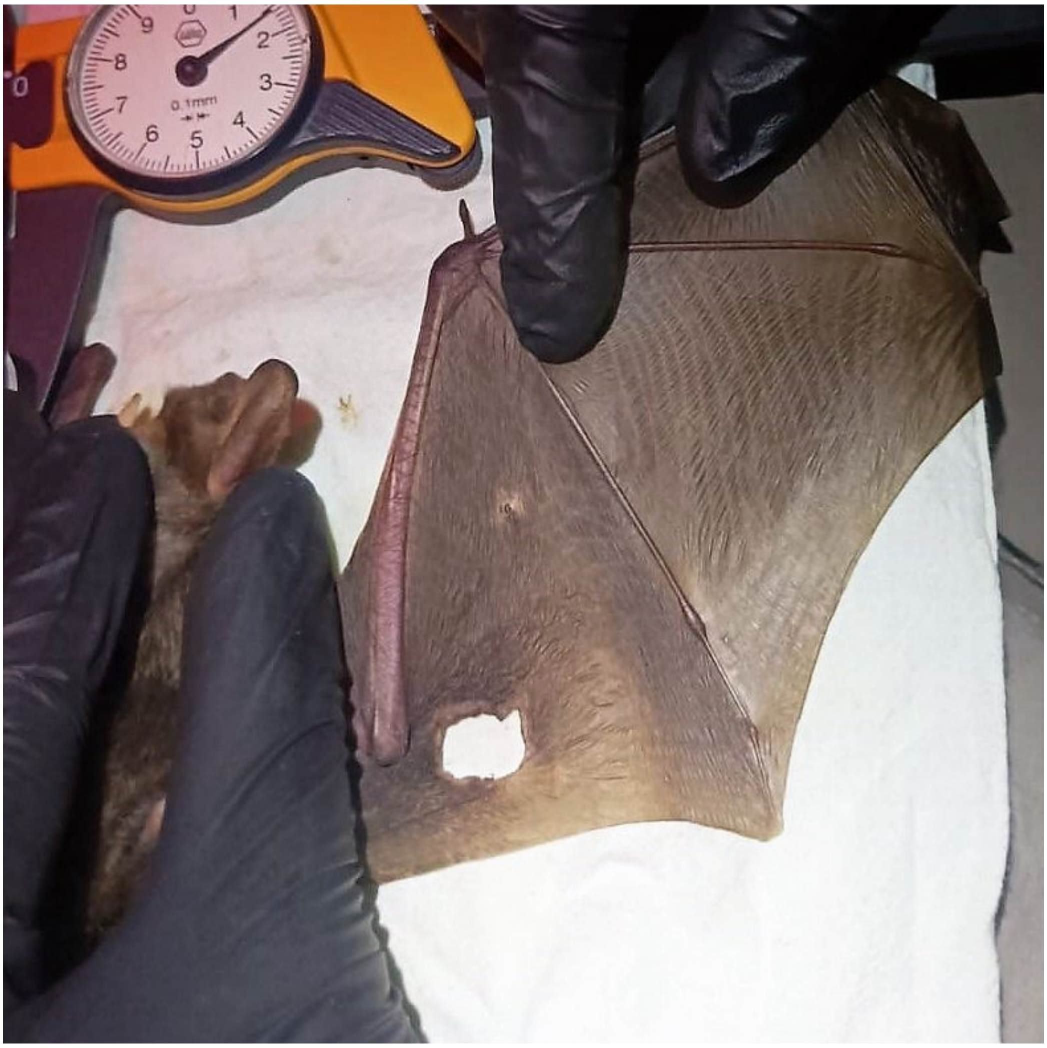

**Figure S10: Wing ruptures of wild-caught greater mouse-eared bats**. Wing-ruptures are often found in Greater mouse-eared bats, which may be caused by collision with gthe ground during gleaning prey captures.

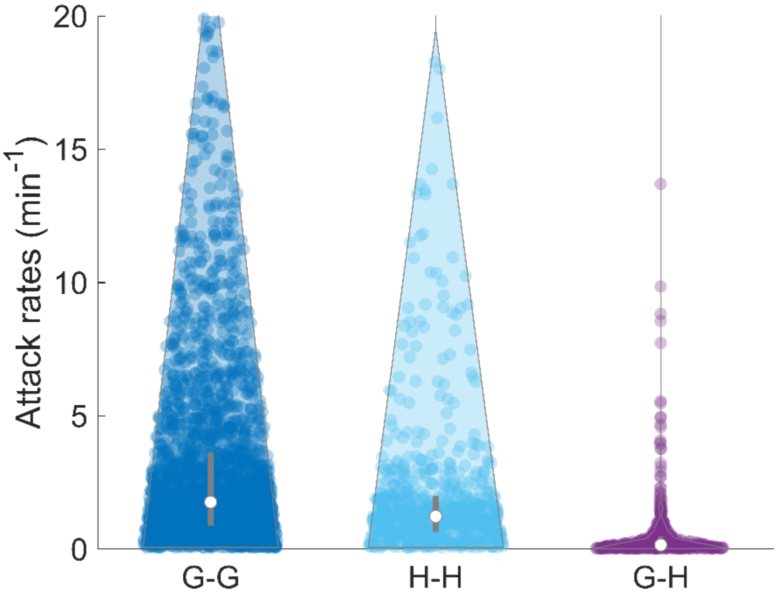


**Figure S11: The bats mainly forage in dedicated foraging bouts.** The attack rate across foraging strategies (purple) are on average 10 times longer than within either gleaning attacks (dark blue) or hawking attacks (light blue).

#### Table S1 Overview of bat individuals, tagging weights and dates

| Bat ID | Tag Type | GPS | Release site | Release Date | Recapture Date | Forearm length (mm) | CM3 | Sex | Status | Bat weight (g) | Tag weight (g) | Tag:Body mass (%) | Recapture weight (g) | Duty cycle (%) |
|---|---|---|---|---|---|---|---|---|---|---|---|---|---|---|
| 1 | A | 0 | OC | 10-07-17 | - | 62.3 | 9.8 | f | PL | 28.6 | 3.5 | 12.2 | - | 100 |
| 2 | A | 0 | DM | 14-07-18 | 17-07-18 | - | - | f | L/PL | 30.1 | 3.7 | 12.3 | 27.3 | 100 |
| 3 | A | 0 | DM | 14-07-18 | 17-07-18 | - | - | f | L/PL | 31.2 | 4 | 12.8 | 29.5 | 100 |
| 4 | A | 0 | DM | 22-07-18 | 24-07-18 | 64 | 9.8 | f | L/PL | 29.8 | 3.8 | 12.8 | 27.1 | 100 |
| 5 | A | 0 | DM | 22-07-18 | 24-07-18 | 67 | 10 | f | L/PL | 30.5 | 4 | 13.1 | 28 | 100 |
| 6 | A | 0 | DM | 22-07-18 | 24-07-18 | 64 | 10 | f | L/PL | 30.1 | 3.5 | 11.6 | 27.1 | 100 |
| 7 | A | 0 | DM | 22-07-18 | 24-07-18 | 64 | 10 | f | L/PL | 29.4 | 3.6 | 12.2 | 25.1 | 100 |
| 8 | B | 0 | OC | 22-07-18 | 23-07-18 | 63.9 | 10 | f | L | 30.1 | 4.3 | 14.3 | 25.8 | 50 |
| 9 | B | 0 | OC | 22-07-18 | 23-07-18 | 63.7 | 10 | f | L | 30.6 | 4.6 | 15.0 | 27.3 | 50 |
| 10 | B | 0 | OC | 22-07-18 | 23-07-18 | 63.3 | 10 | f | L | 31.5 | 4.6 | 14.6 | 26.9 | 50 |
| 11 | B | 0 | OC | 22-07-18 | 23-07-18 | 65.9 | 10.1 | f | L | 30.2 | 4.5 | 14.9 | 26.2 | 50 |
| 12 | B | 0 | OC | 22-07-18 | 23-07-18 | 65.4 | 9.8 | f | L | 31 | 4.6 | 14.8 | 27.3 | 50 |
| 13 | A | 0 | DM | 04-08-18 | - | 66.1 | 10 | f | PL | 29.3 | 4 | 13.7 | - | 100 |
| 14 | B | 0 | DM | 04-08-18 | 07-08-18 | 63.5 | 9.9 | f | PL | 30.2 | 4.6 | 15.2 | 28.8 | 100 |
| 15 | B | 0 | DM | 04-08-18 | 07-08-18 | 66.2 | 10.5 | f | PL | 28.4 | 4.6 | 16.2 | - | 100 |
| 16 | B | 0 | DM | 11-08-18 | 16-08-18 | 65.5 | 10.2 | f | PL | 29.8 | 4.6 | 15.4 | 27.7 | 100 |
| 17 | A | 0 | DM | 11-08-18 | 13-08-18 | 64 | 10 | f | PL | 28.7 | 4 | 13.9 | - | 100 |
| 18 | B | 0 | DM | 11-08-18 | 19-08-18 | 61.2 | 10.2 | f | PL | 29.2 | 4.5 | 15.4 | 27.6 | 100 |
| 19 | A | 0 | OC | 10-07-19 | 12-07-19 | 64.3 | 10 | f | PL | 29 | 3.5 | #DIV/0! | 27.4 | 100 |
| 20 | A | 0 | OC | 10-07-19 | 12-07-19 | 62.3 | 10 | f | PL | 29.3 | 3.5 | 11.9 | 26.9 | 100 |
| 21 | A | 0 | OC | 10-07-19 | - | 66.9 | 10 | f | PL | 29.4 | 3.5 | 11.9 | - | 100 |
| 22 | B | 1 | OC | 11-07-19 | - | 64.5 | 9.9 | f | L/PL | 29.9 | 4.4 | 14.7 | - | 100 |
| 23 | B | 1 | OC | 11-07-19 | 18-07-19 | 65.3 | 10 | f | L/PL | 30.6 | 4.5 | 14.7 | - | 100 |
| 24 | A | 0 | DM | 18-07-19 | 20-07-19 | 63.4 | 9.9 | f | PL | 27.9 | 3.5 | 12.5 | 29.4 | 100 |
| 25 | A | 0 | DM | 18-07-19 | 20-07-19 | 63.4 | 10 | f | PL | 28 | 3.5 | 12.5 | 27.1 | 100 |
| 26 | B | 1 | DM | 18-07-19 | 20-07-19 | 65.3 | 10.2 | f | L/PL | 29.3 | 4.5 | 15.4 | 27.9 | 100 |
| 28 | A | 0 | DM | 18-07-19 | - | 64.1 | 10.1 | f | PL | 28.1 | 3.5 | 12.5 | - | 100 |
| 28 | B | 1 | DM | 18-07-19 | - | 64.5 | 10.1 | f | L/PL | 30.4 | 4.5 | 14.8 | - | 100 |
| 29 | B | 1 | Tabachka | 19-07-19 | 20-07-19 | 63.5 | 10.2 | f | L/PL | 29.4 | 4.5 | 15.3 | 27.8 | 100 |
| 39 | A | 0 | DM | 22-07-19 | - | - | - | f | PL | 28.9 | 3.8 | 13.1 | - | 100 |
| 31 | B | 1 | DM | 22-07-19 | 25-07-19 | 65.6 | 10.1 | f | L/PL | 30.1 | 4.4 | 14.6 | 28.2 | 100 |
| 32 | B | 0 | DM | 22-07-19 | 24-07-19 | 63 | 9.7 | f | L/PL | 29.8 | 4.3 | 14.4 | 28.1 | 100 |
| 33 | B | 1 | DM | 22-07-19 | 25-07-19 | 64.9 | 9.8 | f | L/PL | 29.5 | 4.4 | 14.9 | 28.1 | 100 |
| 34 | B | 0 | DM | 22-07-19 | 25-07-19 | 63.6 | 10.1 | f | L/PL | 29.7 | 4.4 | 14.8 | 28.4 | 100 |

#### Table S2: Capture attempts and success rates for bats trained to forage in a flight room

| **Strategy** | **Aerial** | | | **Ground** | |
|---|---|---|---|---|---|
| | Tag | | No Tag | Bowl | Natural |
| **(N = 2-3 bats)** | Moths | Mealworms | | Small & large beetles | |
| Trials | 116 | 81 | 89 | 56 | 67 |
| Success rates | 0.69 | 0.95 | 0.75 | 0.85 | 0.39 |

**Table S3 Summary statistics on the effect of foraging strategy on the capture attempts (Model 1a, N = 29 bats)**

| ***Model 1a*** | **Capture attempts ~ Strategy + ( 1 \| AnimalNo)** | | |
|---|---|---|---|
| **Distribution** | Poisson (log link) | | |
| | **Fixed effects** | | |
| Parameters | **Est** | **Std. Error** | ***p-value*** |
| *Intercept* | 3.2 | 0.09 | 0 |
| *Strategy* | 1.28 | 0.04 | 0 |
| | **Random effects** | | |
| *AnimalNo* | 0.7 | | |
| | **Goodness of fit** | | |
| $R_{(m)}^2$ | 0.63 | | |
| $R_{(c)}^2$ | 0.96 | | |
| $R^2$ fixed effect | 0.27 | | |

**Table S4 Summary statistics on the effect of foraging strategy on the foraging success (Model 1b, N = 29 bats)**

| ***Model 1b*** | **Success rate ~ Strategy + ( 1 \| AnimalNo)** | | |
|---|---|---|---|
| **Distribution** | Gaussian | | |
| | **Fixed effects** | | |
| Parameters | **Est** | **Std. Error** | ***p-value*** |
| *Intercept* | 32.1 | 3.12 | 0 |
| *Strategy* | 49.22 | 3.8 | 0 |
| | **Random effects** | | |
| *AnimalNo* | 0.08 | | |
| | **Goodness of fit** | | |
| $R_{(m)}^2$ | 0.68 | | |
| $R_{(c)}^2$ | 0.77 | | |
| $R^2$ fixed effect | 0.68 | | |

**Table S5 Summary statistics on the effect of habitat use, foraging strategy and movement style on capture attempts (Model 2a, N = 7 GPS tagged bats)**

Predictor variables: Habitat: Field (0) vs forest (1), Strategy: Gleaning (0) vs hawking (1), and Style: Actively searching for prey (0) vs travelling (1).

| ***Model 2a*** | **Capture attempts ~ Habitat + Strategy + Style + ( 1 \| AnimalNo)** | | |
|---|---|---|---|
| **Distribution** | Poisson (log-link) | | |
| | **Fixed effects** | | |
| Parameters | **Est** | **Std. Error** | ***p-value*** |
| *Intercept* | 2.5 | 0.45 | 0 |
| *Habitat1* | -0.75 | 0.18 | 0 |
| *Strategy1* | -0.93 | 0.20 | 0 |
| *Style1* | 2.02 | 0.23 | 0 |
| | **Random effects** | | |
| *AnimalNo* | 0.78 | | |
| | **Goodness of fit** | | |
| $R_{(m)}^2$ | 0.44 | | |
| $R_{(c)}^2$ | 0.89 | | |

**Table S6 Summary statistics on the effect of habitat use, foraging strategy and style of movement (Model 2b, N = 7 GPS tagged bats)**

| ***Model 2b*** | **Success Rate ~ Habitat + Strategy + Style + ( 1 \| AnimalNo)** | | |
|---|---|---|---|
| **Distribution** | Gaussian | | |
| | **Fixed effects** | | |
| Parameters | **Est** | **Std. Error** | ***p-value*** |
| *Intercept* | 36.8 % | 9.5 | 0.0005 |
| *Habitat1* | -17.9 % | 7.2 | 0.019 |
| *Strategy1* | +67.1 % | 7.7 | 0 |
| *Style1* | -5.0 % | 9.3 | 0.59 |
| | **Random effects** | | |
| *AnimalNo* | 0.0021 | | |
| | **Goodness of fit** | | |
| $R_{(m)}^2$ | 0.72 | | |
| $R_{(c)}^2$ | 0.72 | | |

**Table S7 Summary statistics of the effect of tagging date on the release site and foraging strategy ratio (gleaning:aerial, N = 34 bats)**

| ***Model 3*** | **Strategy Ratio ~ Release Date + (1\|Release Site)** | | |
|---|---|---|---|
| **Distribution** | Gaussian | | |
| | **Fixed effects** | | |
| Parameters | **Est** | **Std. Error** | ***p-value*** |
| *Intercept* | 83.4 | 9.07 | <0.0001 |
| *Release Date2* | -3.1 | 11.1 | 0.777 |
| *Release Date3* | -73.3 | 9.3 | <0.0001 |
| *Release Date4* | 12.2 | 10.6 | 0.26 |
| *Release Date5* | 16.1 | 10.6 | 0.1 |
| *Release Date6* | 15.1 | 10.1 | 0.14 |
| *Release Date7* | 13.1 | 10.6 | 0.2 |
| *Release Date8* | 5.2 | 10.1 | 0.6 |
| *Release Date9* | -23.2 | 12.7 | 0.08 |
| *Release Date10* | -29.6 | 10.1 | 0.007 |
| | **Random effects** | | |
| *Release Site* | 2.93 | | |
| | **Goodness of fit** | | |
| $R_{(m)}^2$ | | | |
| $R_{(c)}^2$ | - | | |
| | **Anova-test of model 3** | | |
| | **DF** | **F-value** | ***p-value*** |
| Intercept | 23 | 589.8 | <0.0001 |
| Release Date | 23 | 62.8 | <0.0001 |

**Table S8: Confusion matrixes of the mastication sound detector**

| | | **Actual** | |
|---|---|---|---|
| **Predicted** | | + | - |
| | **Y** | **TP** | **FP** |
| | **N** | **FN** | **TN** |

| **G** | | **Listening** | |
|---|---|---|---|
| **Detector** | | + | - |
| | **Y** | **132** | **2** |
| | **N** | **4** | **314** |

| **A** | | **Listening** | |
|---|---|---|---|
| **Detector** | | + | - |
| | **Y** | **271** | **3** |
| | **N** | **27** | **86** |

Confusion matrix (left) of gleaning (middle) and aerial (right) prey encounters. Here, TP is true positives, FP false positives, FN false negatives, and TN true negatives.

Actual values were based on the manual listening for mastication sounds, while predicted values were based on the mastication detector. Using the derivatives of the confusion matrix, two metrics were used to describe the performance of the chewing detector:

**positive predictive value (PPV)** = TP/(TP+FP) (1)

**false negative rate (FNR)** = FN/(FN+TP) (2)

| | **Gleaning** | **Hawking** |
|---|---|---|
| **PPV** | **0.99** | **0.99** |
| **FNR** | **0.03** | **0.09** |

**(N = 9 bats)**
